# Generating High Density, Low Cost Genotype Data in Soybean [*Glycine max* (L.) Merr.]

**DOI:** 10.1101/547711

**Authors:** Mary M. Happ, Haichuan Wang, George L. Graef, David L. Hyten

## Abstract

Obtaining genome-wide genotype information for millions of SNPs in soybean [*Glycine max* (L.) Merr.] often involves completely resequencing a line at 5X or greater coverage. Currently, hundreds of soybean lines have been resequenced at high depth levels with their data deposited in the NCBI short read achieve. This publicly available dataset may be leveraged as an imputation reference panel in combination with skim (low coverage) sequencing of new soybean genotypes to economically obtain high-density SNP information. Ninety-nine soybean lines resequenced at an average of 17.1X were used to generate a reference panel, with over 10 million SNPs called using GATK’s Haplotype Caller tool. Whole genome resequencing at approximately 1X depth was performed on 114 previously ungenotyped experimental soybean lines. Coverages down to 0.1X were analyzed by randomly subsetting raw reads from the original 1X sequence data. SNPs discovered in the reference panel were genotyped in the experimental lines after aligning to the soybean reference genome, and missing markers imputed using Beagle 4.1. Sequencing depth of the experimental lines could be reduced to 0.3X while still retaining an accuracy of 97.8%. Accuracy was inversely related to minor allele frequency, and highly correlated with marker linkage disequilibrium. The high accuracy of skim sequencing combined with imputation provides a low cost method for obtaining dense genotypic information that can be used for various genomics applications in soybean.

## Introduction

Genomics research has yielded a variety of tools which allow for more efficient and precise translation of genetic variation into crop improvements. Panels of single nucleotide polymorphisms (SNPs) obtained through SNP arrays or genotyping-by-sequencing (GBS) are the most common tool used to explore and make associations between genetic and phenotypic variation. Genomics-assisted crop breeding continues to demand increasing densities of genotype information to successfully dissect and predict genetically complex traits (Hamblin, Buckler, and Jannink 2011; Lorenz et al. 2011). Current approaches of directly ascertaining a high density of SNP genotype data on large populations are cost prohibitive or fall short of being able capture the maximum amount of genetic space.

Fixed SNP arrays and GBS are popular options for SNP genotyping in crops. Panels ranging in densities of up to ∼600,000 variants are now common in several crop species (Rasheed et al. 2017). However, recent genomics studies are utilizing datasets consisting of one million or more markers to answer complex, quantitative genetic questions. The need for this high density of markers is rendering current arrays and GBS approaches inadequate to generate the magnitude of data modern genomic studies require (Patil et al. 2016; Li et al. 2018; Tian et al. 2011). High-depth whole genome sequencing can achieve these marker densities. One study utilizing high-depth whole genome sequencing in soybean found 9,107,000 high quality SNPs (Valliyodan et al. 2016). Despite advances and the plummeting cost of next generation sequencing (NGS) data, this approach still presents a heavy financial burden, as several reads are required at each variant site to ensure data quality and completeness.

Decreasing genome coverage in the interest of cost savings introduces missing data, which decreases power and can produce biased results. Imputation of missing data has the potential to allow the researcher to recover nearly all of the missing data points resulting from skim sequencing, drastically reducing genotyping expenses associated generating complete, high quality, high resolution SNP datasets. By predicting the unobserved genotypes based on the surrounding variants and their correlation to a complete reference panel, missing data can be amended to the correct allele genotype. This technique has been developed and extensively used in human genomic research, and is now commonly extended to other organisms (Pei et al. 2008; B. N. Howie, Donnelly, and Marchini 2009; B. Howie, Marchini, and Stephens 2011). Seen frequently in plants is the use of imputation to fill missing data points in GBS data (Chung et al. 2017; Chan, Hamblin, and Jannink 2016). Specially designed populations such as bi-parental, nested, and multi-parent where the founders are genotyped to a high depth and used for the reference haplotypes has been shown to boost accuracy (Bayer et al. 2015; Swarts et al. 2014; Tian et al. 2011; Emma Huang et al. 2014; Cericola et al. 2018).

Crop breeding programs working with inbred species and/or inbred lines are uniquely positioned to leverage imputation algorithms in an extremely accurate manner. Near complete homozygosity through inbreeding or double haploids allows calling of genotypes despite having sampled one allele at the site. Large haplotype blocks in historically inbred crops theoretically permit imputation accuracy to extend across large physical regions, where genotyped markers are sparse but in high correlation with each other. Success with such a combinatorial approach has been reported in rice, using ∼1X coverage sequence data of 517 individuals. Imputation of the missing genotypes in these individuals without a reference panel to produce a SNP panel of ∼3.6 million markers with > 98% accuracy (X. Huang et al. 2010). This was confirmed in a later study that also included simulations performed down to 0.1X depth. Falling below a depth of 0.5X resulted in steep accuracy consequences, with concordance falling to 76% at the 0.1X level. (Wang et al. 2016).

Incorporation of a reference panel has been shown to result in large accuracy improvements at sequencing coverage less than <1X in humans, where imputation at the 0.1X level was improved from less than 5% accuracy to ∼70% (Pasaniuc et al. 2012). With the growing amount of sequence data present in public databases for many common crops, it is possible to generate an extensive reference panel that might improve accuracy at ultra-low sequence coverage and further cut per sample genotyping cost. In this study, we report on a low coverage whole genome sequencing with imputation approach in a naturally inbred crop, soybean, for producing a low cost, high quality, high density SNP dataset. A reference panel was generated using publicly available high-depth sequencing data for 106 lines, and employed for imputing the missing genotypes of 114 lines sequenced at ultra-low depth. Coverages from 0.1X – 1X depth at intervals of 0.1X were evaluated. The factors influencing error rates and extensibility within/outside soybean were investigated, and the consequences of error rates and types of error on a typical genome-wide association study (GWAS) were explored.

## Materials & Methods

### Reference Panel

The reference panel for genotype imputation was generated using publicly available sequence data deposited in the NCBI Short Read Archive from study number SRP062245 (Valliyodan et al. 2016). This unfiltered, raw dataset consisted of 106 *Glycine max* lines sequenced at an average of 17.1X coverage **(Supplementary Table 1)**. The raw reads were filtered for adapter sequence contamination, base quality, and truncated reads using Trimmomatic (Bolger, Lohse, and Usadel 2014). Bowtie2 was used to map reads to the *Glycine max* Wm82.a2.v1 reference genome with the “very sensitive” option (Langmead and Salzberg 2012). Reads with a mapping quality score of less than 20 were discarded. SNPs were called using the GATK3.7 HaplotypeCaller tool for an initial panel of 13,052,759 SNPs across all lines (Poplin et al. 2017). SNP calls with five or less reads supporting the call were filtered out, as well as calls with a confidence score of less than 20. To control for potential sample contamination/mixing, the inbreeding coefficient, also called the F statistic (JAIN and WORKMAN 1967), was calculated using the software Plink1.9 (Shaun Purcell et al. 2007). As soybean is historically an inbred crop, one can expect F statistics close to one in *Glycine max*. Seven samples fell below a cutoff of 0.9 and were discarded from the final reference panel. All heterozygous calls in the remaining 99 lines were filtered, leaving only biallelic SNPs for consideration. The final reference panel spanned 10,803,148 biallelic homozygous SNPs in 99 lines compared to 10,417,285 SNPs found by Valliyodan et al using the same data set.

**Table 1:** The number of markers and genotyping rate in each low coverage subset from 0.1X to 1X sequencing depth. As coverage decreases, the total number of markers captured and completeness of the SNP panel decreases.

| Mean Depth | Genotyping Rate | Number of SNPs | Reads | Base Pairs |
| --- | --- | --- | --- | --- |
| 1 | 32.44% | 1,288,463 | 6,327,889 | 949,183,385 |
| 0.9 | 30.41% | 1,240,823 | 5,695,100 | 854,265,047 |
| 0.8 | 27.77% | 1,174,619 | 5,062,311 | 759,346,708 |
| 0.7 | 24.91% | 1,097,843 | 4,429,522 | 664,428,370 |
| 0.6 | 21.80% | 1,005,880 | 3,796,734 | 569,510,031 |
| 0.5 | 18.47% | 895,596 | 3,163,945 | 474,591,693 |
| 0.4 | 14.98% | 760,167 | 2,531,156 | 379,673,354 |
| 0.3 | 11.40% | 590,786 | 1,898,367 | 284,755,016 |
| 0.2 | 7.85% | 375,343 | 1,265,578 | 189,836,677 |
| 0.1 | 4.74% | 133,747 | 632,789 | 94,918,339 |

### Imputation Panel

To generate a low sequence coverage panel for imputation, 114 experimental lines selected form the University of Nebraska soybean breeding program **(Supplementary Table 1)** were sequenced to a depth of 1X or greater on an Illumina NextSeq 500 (Illumina Hayward, Hayward, CA) using the manufacturer’s protocol and 150 base pair paired end reads. DNA was isolated from lyophilized leaf tissue collected from twenty plants per genotype a CTAB method based extraction method (KEIM and P. 1988) scaled down for a 96 well plate by dividing all reagent volumes by 40. Extracted genomic DNA was fragmented using a Covaris S220 with the manufacturer’s recommended settings for generating ∼350 base pair length fragments (Covaris, Inc., Woburn, MA 01801). Double sided size selection was performed using KAPA Pure Beads to retain only fragments within the 250-450 base pair range using the manufacturer’s protocol and eluted in 40 μl of TE buffer (Roche Sequencing Solutions, Santa Clara, CA 95050). After testing DNA concentration, samples were standardized to 62.5 ng /μl. Libraries were prepared using a custom protocol adapted from literature to perform A-tailing and end-repair in one reaction, and avoid PCR after adapter ligation by extending the incubation time (Kozarewa and Turner 2011; Knapp, Stiller, and Meyer 2012). To perform end repair and A-tailing, 16 μl of fragmented genomic DNA for each sample was combined with 1 μl of T4 polynucleotide kinase (PNK) (10U/μl), 1 μl of T4 DNA polymerase (5U/μl), 1 μl of DreamTaq DNA Polymerase (5U/ μl), 2.7 μl of Cut Smart Buffer (10x), 2.2 μl of dATP (10mM), 0.8 μl of dNTP (10 mM), and 0.3 μl of ATP (10mM). Samples were incubated in a thermocycler for 30 minutes at 20°C, and then immediately ramped to 65°C and held at this temperature for 30 minutes. Samples then proceeded immediately to adapter ligation. To the 25 μl of end repaired and A-tailed product the following was added: 10 μl of T4 DNA Ligase Buffer, 3 μl of T4 DNA Ligase (2000U/μl), 3 μl of PEG 6000, 2 μl of PCR grade water, and 2 μl of uniquely barcoded adapters (30mM). Samples were incubated on a thermocycler for 45 minutes at 20°C. After this time, samples were immediately cleaned using KAPA Pure Beads to retain fragments within the 350-550 base pair range and eluted in 20 μl of TE buffer. Multiplexing was performed by combining 5 μl of each individual library. Libraries were quantified using the KAPA Library Quantification Kit for Illumina platforms.

To create subsets simulating depths from 0.1X to 1X at intervals of 0.1X, reads were randomly selected from the raw datasets. Each dataset was trimmed for adapter contamination, base quality and truncated reads using Trimmomatic, and then mapped to the *Glycine max* Wm82.a2.v1 reference genome with Bowtie2 using the “very sensitive” option. Mapped reads below a quality score of 20 were filtered. The genotypes at all 10,803,148 SNP positions in the reference panel were called in the low coverage imputation panel using GATK3.7 Haplotype Caller. Genotyping SNPs from a single read has been found accurate in rice whole genome sequencing and maize GBS applications (Wang et al. 2016; Swarts et al. 2014). Any heterozygous calls were discarded, as well as calls not matching the two allele options at that position. For each subset, a random 5% of calls were masked and considered “true” genotypes for evaluating imputation accuracy.

### Imputation Concordance Evaluation

For the sake of computational efficiency, imputation was performed on a per chromosome basis using Beagle 4.1 (Browning and Browning 2016) with the low memory option. To assess accuracy, the imputed genotype calls were compared to the masked calls, and the percent of those in agreement constituted overall concordance using GATK 3.7’s Genotype Concordance tool (McKenna et al. 2010). This accuracy assessment was performed across sequencing depths and minor allele frequencies. Three post imputation datasets were considered to quantify any accuracy improvement obtained by filtering poorly imputed sites. This included the raw imputed dataset, and two datasets filtered on Beagle’s posterior GP. Values with GP scores under 0.45 and 0.9 were filtered for the latter two evaluation panels, respectively. VCFtools0.01.12a (Danecek et al. 2011) was used to bin by minor allele frequency, and Plink1.9 was used to filter on GP score. GP score filtering thresholds were determined after examining their relationship to error rate (**Supplementary Table 2**)

### Error and Linkage Disequilibrium

Error in relationship to linkage disequilibrium (LD) was examined as a potential metric of extensibility to other soybean population and crop species. D’ and r^2^ statistics were calculated for all pairwise reference panel SNPs using Plink1.9 (Gaunt, Rodríguez, and Day 2007). Proportion of errors made at each SNP site across was calculated by comparing the imputed values to the masked values across all subsets of depths. To reduce noise, data was smoothed through the application of a rolling average window with a width of 1500 SNPs after ordering by the respective LD metric. A second order polynomial was fit to describe the D’ and error relationship, and a simple linear regression was fit to describe the relationship between r^2^ and error.

### Relationship Between Samples & Reference Panel

Close relatedness between the sample and reference genotypes has been previously reported to increase imputation precision. Relatedness matrices were generated based on five different coefficients and averaged the top five scores from each sample genotype as a metric for gauging degree of relatedness to the reference panel. These measures were plotted against concordance scores from the imputed data filtered for GP scores above 0.9 and averaged across all depth levels. A simple linear regression model was fit to assess potential correlation. Relatedness matrices were calculated using the R package “synbreed”, using options corresponding to measures described by vanRaden, Astle and Balding, Reif, Hayes and Goddard, and Euclidean distances (Wimmer et al. 2012).

### Genome Representation

Genomic studies improve as the linkage between the genotyped polymorphism and underlying causative gene increases. The extent of LD between two markers therefore constitutes proxy for the correlation of the marker and underlying gene(s) of interest. To assess how well the panel represented variation across the genome, the distribution of LD in this dataset was compared to the SoySNP50k Array (Song et al. 2015). SNPs with MAF below 0.05 were filtered out, a quality control step implemented in most genomic studies. Both D’ and r^2^ were calculated using Plink1.9, and distributions plotted in R3.4 (Team 2017).

### Error and Beagle Posterior Genotype Probability

To explore the possibility of using the posterior genotype probability (GP) as a post imputation filtering metric, proportion of error across depth subsets was plotted against GP. A rolling average window with a width of 500 SNPs was applied to the proportion error after ordering by GP, and a second degree polynomial was fit to describe the relationship in R3.4.

#### Error Type

Allele frequencies exhibit some degree of influence on the results of many genomics studies. Therefore, how imputation error skews this metric is of significant interest. Masked and imputed datasets were coded according to the major allele in the reference dataset. Errors were binned into four categories, based on which alleles (homozygous major/ homozygous minor/heterzygous) were incorrectly imputed and which allele was true. Because all heterozygous calls were filtered in the initial data generation, no heterozygous to major, or heterozygous to minor category exists.

### Power Analysis

In the interest of determining the potential cost of imputation error, a basic power calculation for minor to major and major to minor errors in a GWAS was performed. Using an R implementation of Purcell’s “Genetic Power Calculator” (S Purcell, Cherny, and Sham 2003), power was calculated to detect a moderate effect QTL across minor allele frequency bins from < 0.025 to 0.5. Simulations assumed an additive genetic model, 300 genotypes, LD between the QTL and marker of 0.8 D’, a significance threshold that mirrored the Bonferroni correction for 1,716,234 SNPs (the final size of the SNP dataset after quality control filtering), and a QTL effect size of 1 standard deviation. Error rates from 1-10% were tested at intervals of 1%. To investigate the possibility of including more genotypes to overcome power losses associated with imputation error, simulations were also performed for 150, 500, and 1000 genotypes for a 5% error rate at the same conditions as specified above.

### Cost Analysis

Decreasing cost per sample allows a researcher to expand a study to overcome power loss introduced through the imputation error. To illustrate the impact of this, per sample sequencing costs were calculated using current Illumina NextSeq500 high throughput 300 cycle sequencing kit prices, cost analysis of a custom library prep protocol, and CTAB DNA extraction method **(Supplementary Table 2)**. The retained cost per sample and average raw concordance were plotted as depth decreased.

### Data Availability

Raw sequencing data directly generated by this project for use in creating the study panel has been deposited in the NCBI Short Read Archive under accession number PRJNA512147. The reference panel used for genotype imputation was generated using previously publicly available sequence data deposited in the NCBI Short Read Archive from study number SRP062245 (Valliyodan et al. 2016).

## Results

### SNP Genotyping & Imputation

The reference panel for imputation was constructed using 106 *Glycine max* lines sequenced at an average of 17.1X coverage using publicly available sequencing data deposited in the NCBI Short Read Archive (Valliyodan et al. 2016) (**Supplementary Table 1)**. After quality control measures were applied to the raw and mapped sequence data (see Materials & Methods), a final reference panel of 10,803,148 biallelic homozygous SNPs across 99 lines was generated. SNPs discovered in the reference panel were used to genotype experimental lines in the study panel. This consisted of 114 lines that were sequenced to a depth of at least 1X. Coverages from 0.1X to 1X were analyzed by randomly subsetting reads from the raw sequence data. Of the 10,803,148 million markers discovered in the reference panel, the number of SNPs genotyped by this low coverage study panel subsets ranged from 133,747 to 1,288,463 markers. These subsets also ranged in missing data rates from 95.26% to 67.56% for those markers **(Table 1)**. Using the reference panel, genotype values for all missing positions were imputed.

An alternative to this whole genome sequencing approach are fixed SNP arrays. However, this method provides less total SNPs for genomic studies and may not capture as much of the genome. High LD between SNPs can be extended to assume a strong correlation to other genomic variation between them. Plotting the density distributions of r^2^ and D’ LD measures for the Soy50KSNP Array and imputed dataset demonstrated that whole genome sequencing with imputation had a greater concentration of values towards higher linkage values. Generally, a D’ or r^2^ of over 0.8 between is considered “strong linkage”. The imputed dataset provided 1,716,234 SNPs after common quality control filters, with 36.00% and 85.66% of r^2^ and D’ values above 0.8, respectively. This is in comparison to the 42,133 SNPs in the fixed array, where 24.20% and 80.00% of r^2^ and D’ values are above 0.8 **(Figure 2)**. If high LD indicates a better tagging of underlying variation, the imputed dataset captures the genome’s SNP variation better than the Soy50KSNP Array.

**Figure 1:**
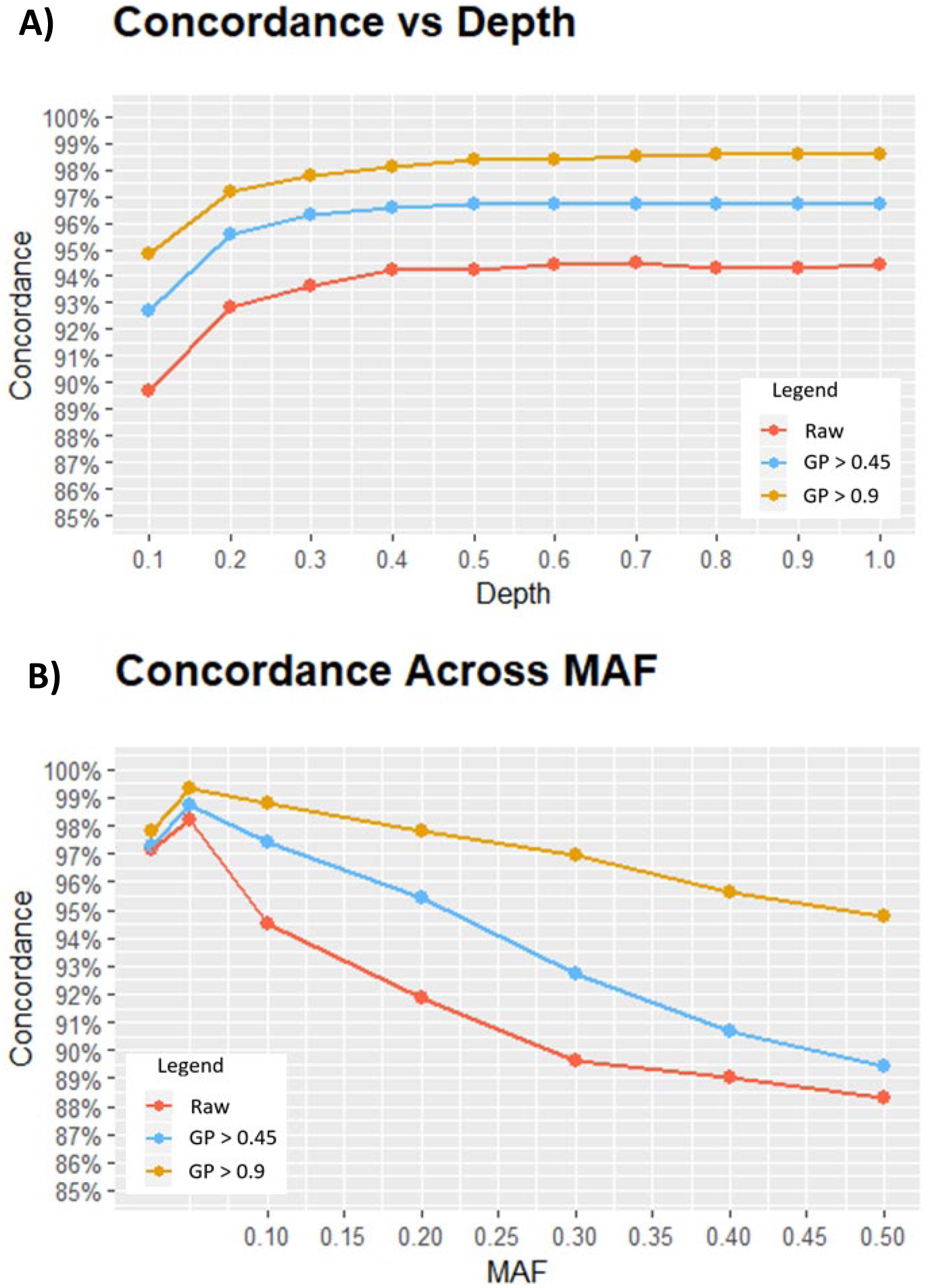
A) Overall accuracy of filtered and raw imputed datasets were plotted across the evaluated depths. For all study panels, concordance rapidly erodes below a sequencing depth of ∼0.3X. B) Examining accuracy in the context of minor allele frequency reveals that error occurs at higher rates as MAF approaches a maximum of 0.5.

**Figure 2:**
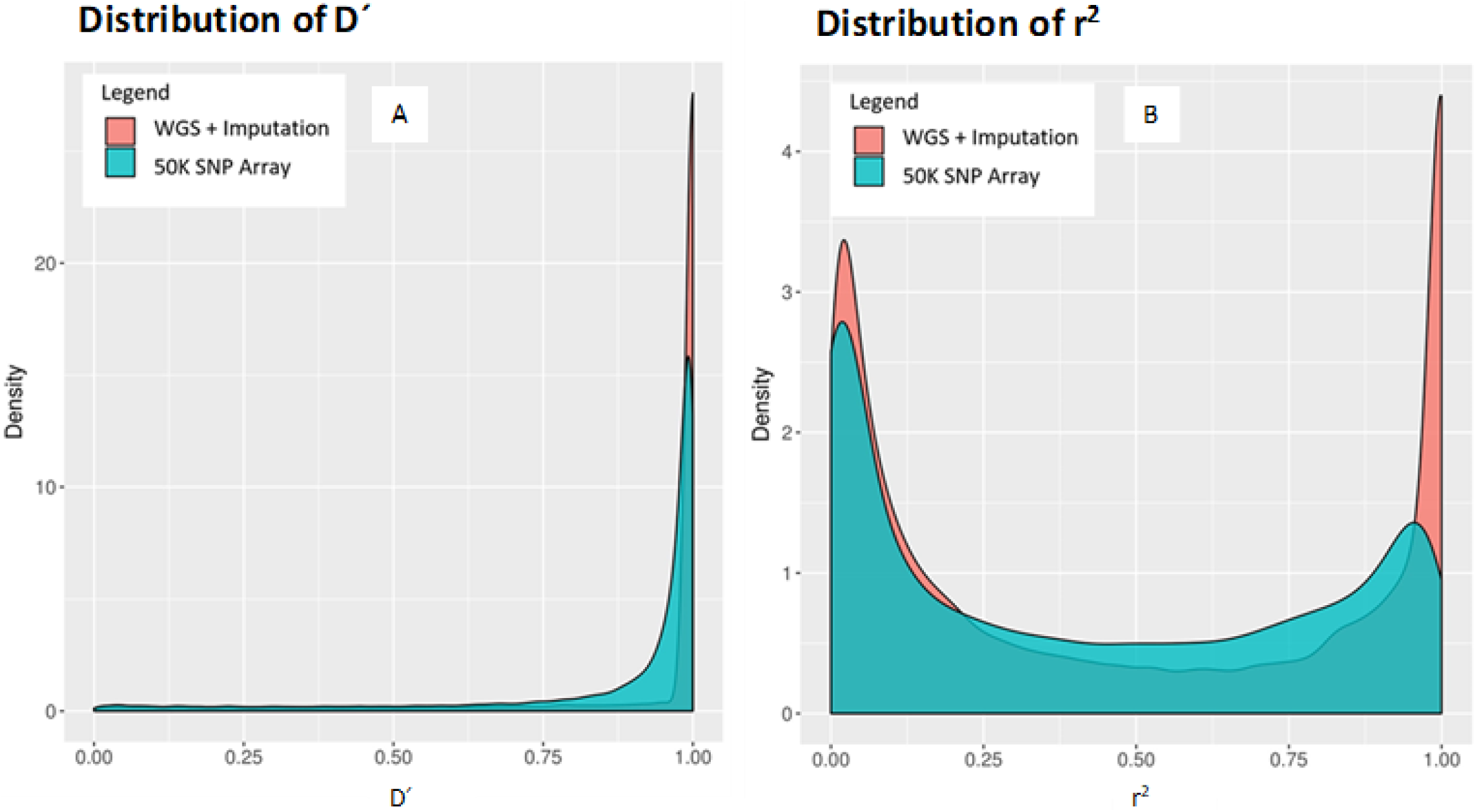
Comparing density plots for LD measures D’ (A) and r^2^ (B) demonstrates that using whole genome sequencing with imputation results in a dataset that has a higher proportion of SNPs is strong pairwise linkage with each other, represented in the heavier tails in red near D’ and r^2^ values of 1.

### Imputation Accuracy

Prior to imputation, 5% of genotype calls from the skim sequencing data were withheld to assess accuracy. Overall imputation accuracy was consistent for raw and filtered datasets as sequencing depth decreased from 1X until 0.3X, where accuracy drops off by an average of 3.5% from 0.3X to 0.1X **(Figure 1A, Supplementary Table 2)**. Assessing the error type of this study showed that 53.13% of the errors made were incorrect imputation of the minor allele when the major allele was true. Of the remaining errors, 35.10% were incorrect imputation of the major allele when the minor allele was true, and 11.77% were incorrect imputation of heterozygous calls. No heterozygous to major/minor errors exist as at heterozygous calls were filtered in the initial panels **(Figure 3)**.

**Figure 3:**
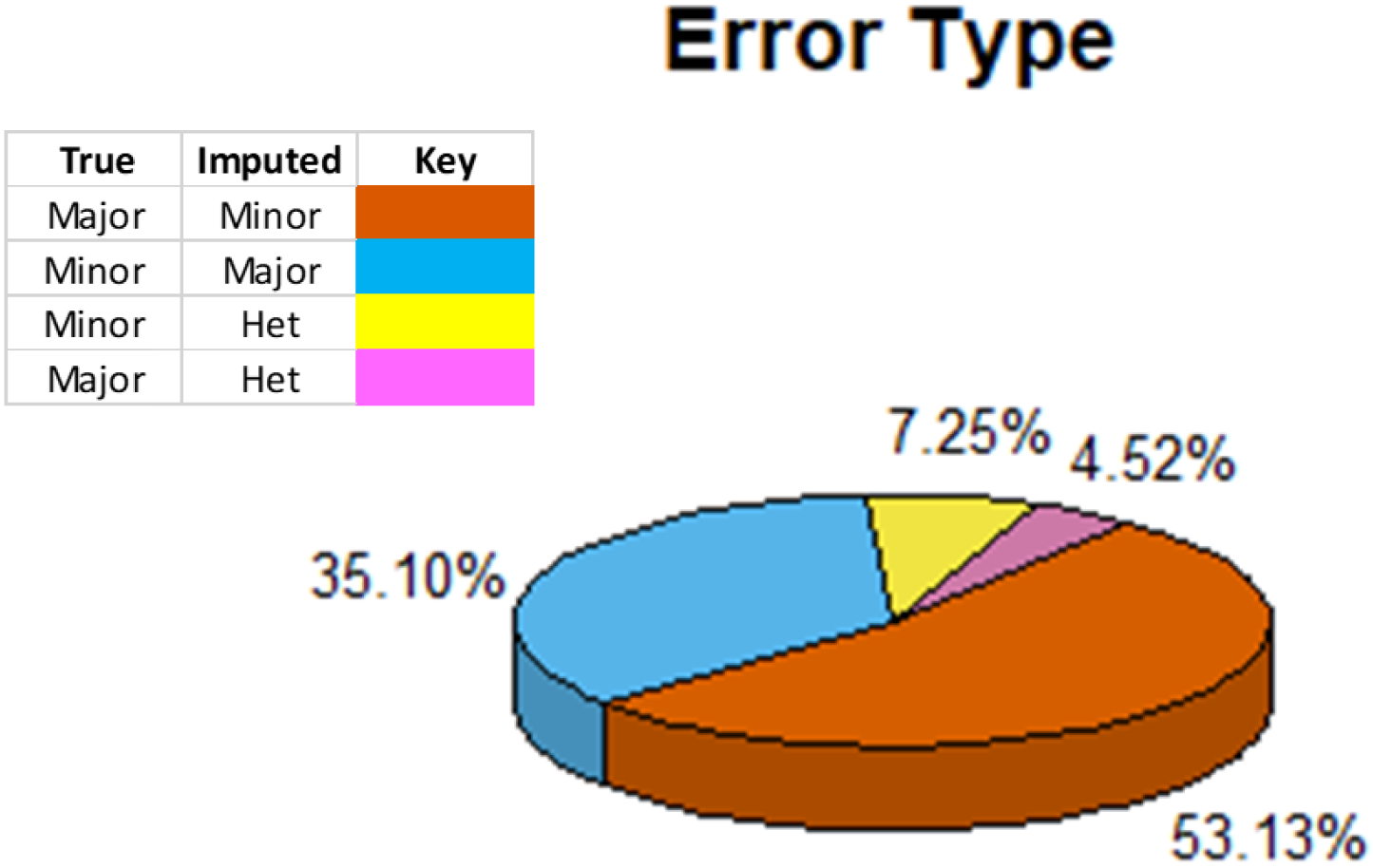
Proportion of errors made as categorized by whether the minor/major/heterozygous alleles was misimputed. In over half of all the errors made, Beagle overimputes the minor allele when the major allele is the true genotype. Incorrect heterozygous imputations make up a minor proportion of the total error and would likely be filtered out in inbred panels.

Filtering on Beagle’s posterior genotype probability (GP) to improve dataset quality was successful. When imputed positions with a GP score of less than 0.45 were discarded, accuracy improved by an average of 2.50% across sequencing depths. A more stringent filter that only kept positions with a GP score over 0.9 resulted in a 4.26% increase in accuracy **(Supplementary Table 2)**. This practice did reintroduce some missing data, which varied across depth and filtering level. Data loss as a result of post imputation filtering was below 5% for all depths at a filtering level of GP > 0.45, but quickly inflates when filtering for imputation quality of GP > 0.9 to a missing data rate of 20.82% at 0.1X **(Supplementary Figure 1).** While filtering on Beagle’s posterior genotype probability may reduce falsely imputed genotypes, it must be balance with the reintroduction of missing data it causes.

The error rate at individual marker loci may not be well captured by the overall concordance across all SNPs. Examining concordance in the context of minor allele frequency (MAF) reveals as MAF values approach a maximum of 0.5, concordance decreases. Application of post imputation filters of GP values increases overall accuracy through improved concordance at these increased MAFs. This trend is uniform across all sequencing depths **(Supplementary Figure 2).** Through examining imputation accuracy in this manner, it is apparent that higher error rates are occurring at SNP positions at MAFs nearest 0.5 than is described by the average concordance measure.

Error rates in imputation may be influenced by characteristics specific to the population and crop species to which it is applied. The correlation between variants is a cornerstone to the success of imputation. If the correlation between alleles is high then imputation accuracy should also be high and as the correlation between alleles decrease then the accuracy of imputation should also decrease. This correlation between alleles can be measured with LD. Soybean is a historically inbred crop with long ranging LD (Zhou et al. 2015). As D’ and r^2^ approach 1, where neighboring SNPs are in perfect linkage with each other, error rates are at their lowest. Both relationships demonstrate a very strong correlation with R^2^ values of 0.98 and 0.89 for r^2^ and D’ respectively **(Figure 4)**, indicating LD is an important factor to consider when applying this technique to other soybean populations or other crop species.

**Figure 4:**
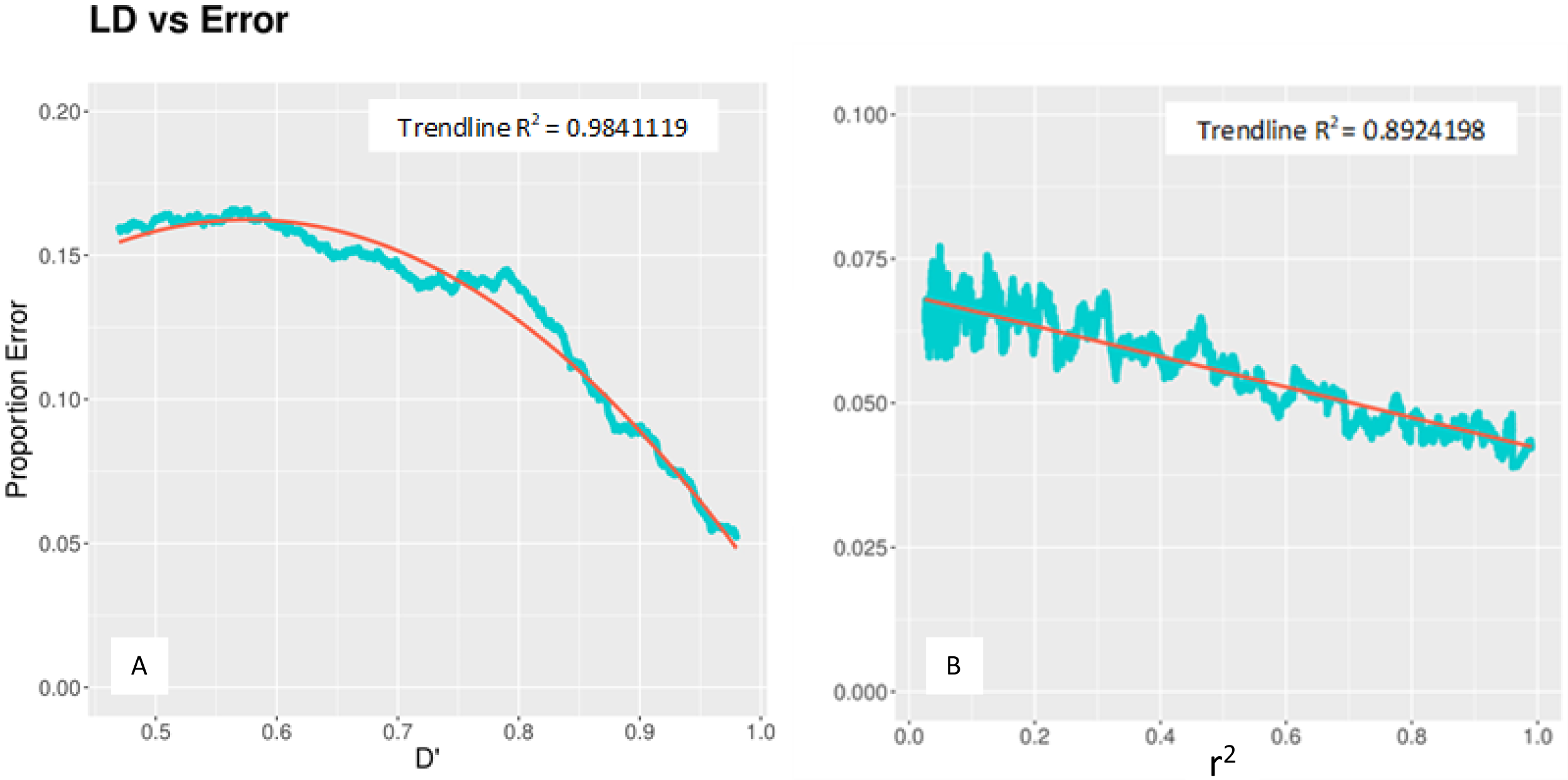
Comparing the smoothed frequency of errors made at individual SNP sites with LD measures D’ (A) and r^2^ (B) demonstrates the strong influence of linkage disequilibrium on imputation accuracy.

Relationship of the study genotypes to the reference panel genotypes has been suggested as a strong influencer of imputation accuracy. Plotting calculated values for five unique kinship metrics against concordance for each genotype did not demonstrate any strong linear relationships. The maximum correlation for any of the measures was for Reif’s method, at an R^2^ of 0.26. Examination of the standard error shows that the study population varies narrowly in terms of relatedness to the reference panel. Additionally, assessing the raw values suggests that the study population is weakly related to reference genotypes. This is best illustrated with the vanRaden and Astle & Balding measurements, where a “strong” relationship is usually indicated by values approximately >= 0.4. In both these cases, the largest measure does not exceed 0.18 and 0.16 (VanRaden 2008; Astle and Balding 2009). The combination of diminished values and narrow standard error indicates a weak relationship of the study panel to the reference panel. **(Supplementary Figure 3)** The evidence of a weak relationship suggests that relationship was not a strong influencer of the high imputation accuracies obtained.

### GWAS Power

Understanding the effect error rate has on genomic studies is important when selecting an appropriate genotyping technique. TO determine the effect of the error rate of skim sequencing and imputation has on GWAS we performed power simulations of detecting a moderate effect QTL in a panel of 300 individuals. This power study showed significantly decreased power to detect QTL with increasing errors at MAF from 0.1-0.3. This was most pronounced when the minor allele was incorrectly imputed as the major allele. Above 0.3 MAF, power for QTL detection was minimally affected by error **(Figure 5A)**. Studying the effect of three additional sample sizes, while assuming a 5% error rate demonstrates the potential for experimenters to recover power losses through inclusion of more genotypes. Including 500 individuals at this fixed error rate recovers and even slightly improves power at mid-range MAFs over studying 300 genotypes with no genotyping error (**Figure 5B)**.

**Figure 5:**
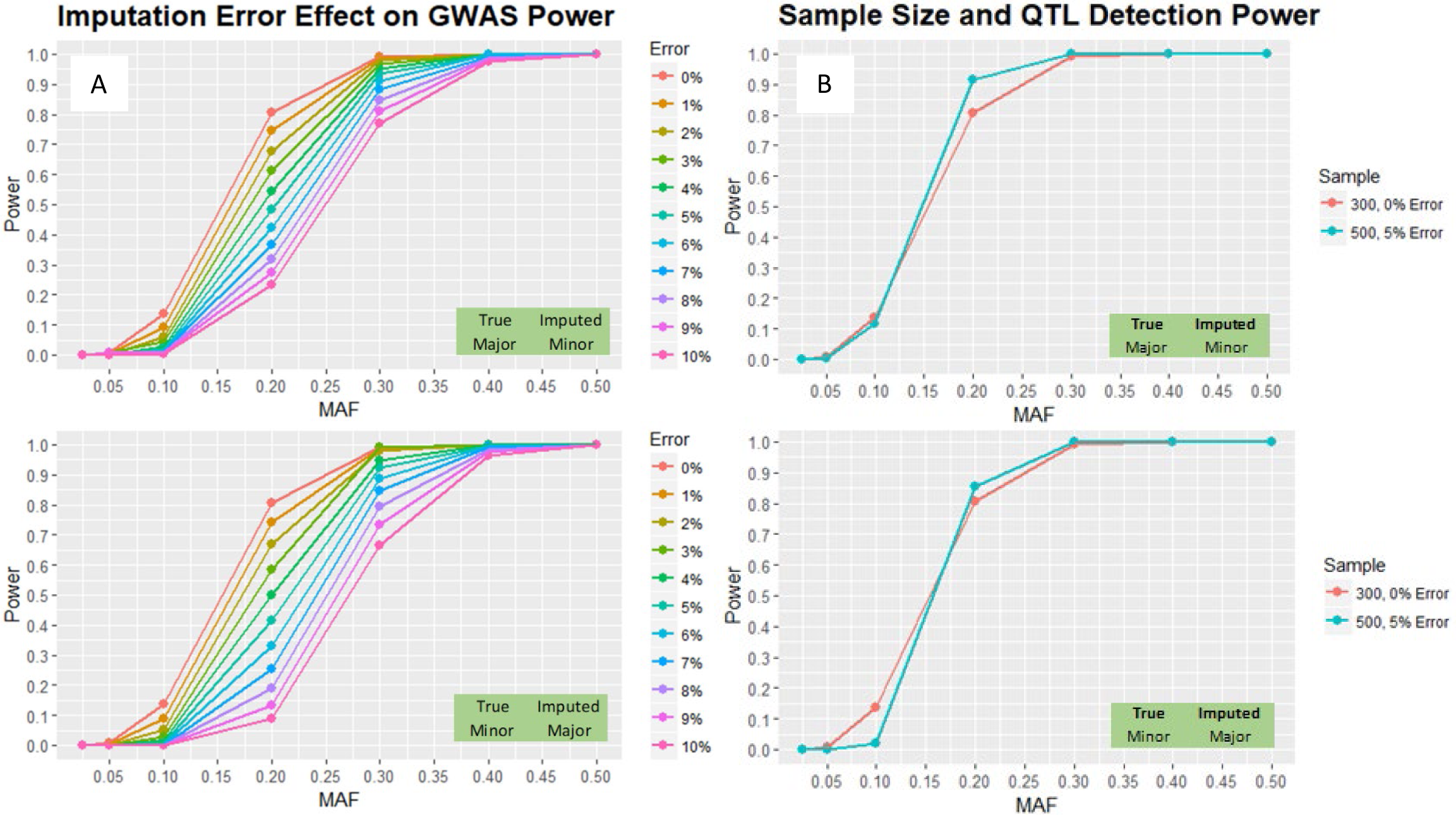
A) The power to detect a moderate effect QTL becomes increasingly sensitive to error for both major to minor and vice versa errors at intermediary MAFs. B) Comparing the power to detect the same QTL with 300 samples at a 0% genotyping error versus 500 samples with a 5% error rate demonstrates that cost savings can be used to increase study sizes in order to recover power losses introduced by the imputation error of both major and minor alleles.

## Discussion

This study illustrates the potential of low coverage sequencing with imputation as an economical approach to obtaining high density SNP genotype information in soybean. Accelerating improvement of complex phenotypes through genomics necessitates high quality, high resolution marker data. However, studies are often limited by the cost required to obtain this information through high coverage sequencing. The combination of low coverage sequencing with imputation presents an option that drastically cuts costs while retaining a high level of accuracy. Implementing a similar method in rice allowed researchers to generate a high quality, dense SNP dataset using 1X depth whole genome sequence (X. Huang et al. 2010; Wang et al. 2016). This analysis in soybean, which differs in the inclusion of a reference panel for imputation, determined sequencing depth could be reduced to 0.3X with no significant accuracy losses. Analogous results have been demonstrated in humans, where it was concluded that a reasonably accurate and dense dataset could be obtained from 0.2X coverage supplemented with imputation using a reference panel (Pasaniuc et al. 2012). To our knowledge, this is the first work to examine using imputation with real sequence data at less than 1X coverage in the construction of a high quality, highly affordable SNP dataset in plants. The effect of imputation method and structure of the reference panel have not been specifically examined in the context of application to skim sequencing, providing future avenues for research and improvement.

While SNP arrays and GBS are popular options for obtaining genotype information, high precision genomics demands markers to be in close linkage to the contributing genes. Regions of the genome with sparsely correlated markers may therefore contain overlooked causal variation (Hirschhorn and Daly 2005; Witte 2010). Skim sequencing with imputation, as investigated here, tags a significantly larger portion of the genome in tighter LD than the current soybean 50k array. This effect may be presumed to extend to GBS datasets of a similar size. Such a boost in resolution may therefore reveal QTL in regions of the genome that would not have been captured through smaller datasets.

The accuracy and extensibility of this approach in other soybean populations, as well as other crops is based on several factors. To explore potential limitations in this method, population LD and the relatedness of reference panel to study lines were examined. Both of these factors have been implicated as strong influencers of imputation accuracy due to the innate reliance of the technique on the presence of sample haplotypes within the reference panel, as well as the extent of correlation between observed markers (Hickey et al. 2012; He et al. 2015). The strong inverse relationship observed between the proportion of SNPs incorrectly imputed at a given position and LD measurements suggests that for soybean populations and other crops with shorter range LD, imputation accuracy will likely decrease. There was no significant relationship detected between kinship measures and accuracy. However, the study genotypes exhibited little variation for any of the calculated metrics, which can be seen in the low standard deviations. Without a wide range of values to examine, identifying a clear trend is unlikely. The positive effect of relatedness on imputation accuracy is documented in other literature (Hickey et al. 2012; Ma et al. 2013; Boison et al. 2015), and should therefore be a consideration in expanding this method to other soybean populations and crop species. The overall weak kinship between study and reference panels in this data may also be viewed as a positive, since high levels of imputation accuracy were achieved despite this populations being interpreted as distally related.

The power to detect a QTL is partially dependent on the allele frequency at that loci (Ardlie, Lunetta, and Seielstad 2002; Tabangin, Woo, and Martin 2009). Therefore, the relationship between imputation accuracy, minor allele frequency (MAF), and statistical power may be considered particularly important. In agreement with an analysis performed with maize, the data showed steadily decreasing imputation accuracy as MAF increased with the exception of very rare alleles (MAF < 0.05) (Hickey et al. 2012). An opposite tendency was observed with respect to statistical power losses across MAF, so it can be interpreted that at the loci a SNP dataset would display the highest imputation error rates, the GWAS is least affected by them. This trend has also been supported in human imputation analyses looking at sample size inflation factors under different imputation error types (L. Huang, Wang, and Rosenberg 2009). In both cases, power consequences were greater for incorrect imputation of the minor allele. It is unclear how a combination of error types at a SNP locus would influence genomic studies. Decisions on the level of decreased coverage that can be tolerated should consequently be made not on the overall average concordance, but by examining the concordance across minor allele frequencies in relation to the maximum allowable error to retain power.

The cost savings associated with this method can be used to include more sample genotypes, not only recovering power losses at low minor allele frequencies, but potentially increasing total power. Similar results in humans have indicated sampling more genotypes with small error is more beneficial over fewer genotypes with perfect accuracy (Pasaniuc et al. 2012). Comparing the raw accuracy along sequencing depths along with per sample costs, shows that at the previously identified critical threshold of 0.3X coverage, there is only a 0.85% loss of accuracy relative to using a 1X sequence, while costs decreased 57% **(Supplementary Figure 4)**. Moreover, the use of public sequence data to construct a broad reference panel eliminates the cost and limitations of assembling special populations and sequencing the founders to a high coverage to serve as the reference haplotypes.

## Conclusion

Here it is demonstrated that low coverage sequencing accompanied with imputation from a reference panel can be extended below 1X depth in soybean to capture high density, reasonably accurate SNP genotype information economically. The tremendous drop in per sample sequencing cost over high depth methods may allow researchers to expand the number of study genotypes in their investigations, while representing a larger portion of the genome than fixed SNP arrays and GBS data. The potential for success of this genotyping method within and outside of soybean is highly reliant on population LD. Furthermore, researchers should examine accuracy and power within the context of minor allele frequency to make informed decisions about sequencing depth tolerances. As genomics demands increasing SNP panel densities across a wide range of genotypes, skim sequencing with imputation constitutes a financially feasible and highly accurate way to meet these requirements.

## Acknowledgements

Research reported in this publication was supported by the Nebraska Soybean Board project #1726. The authors would also like to acknowledge Dr. Reka Howard and Dr. Keenan Amundsen for providing their technical perspective during the compilation of project results and manuscript drafting.

---

**Supplementary Figure 1:**
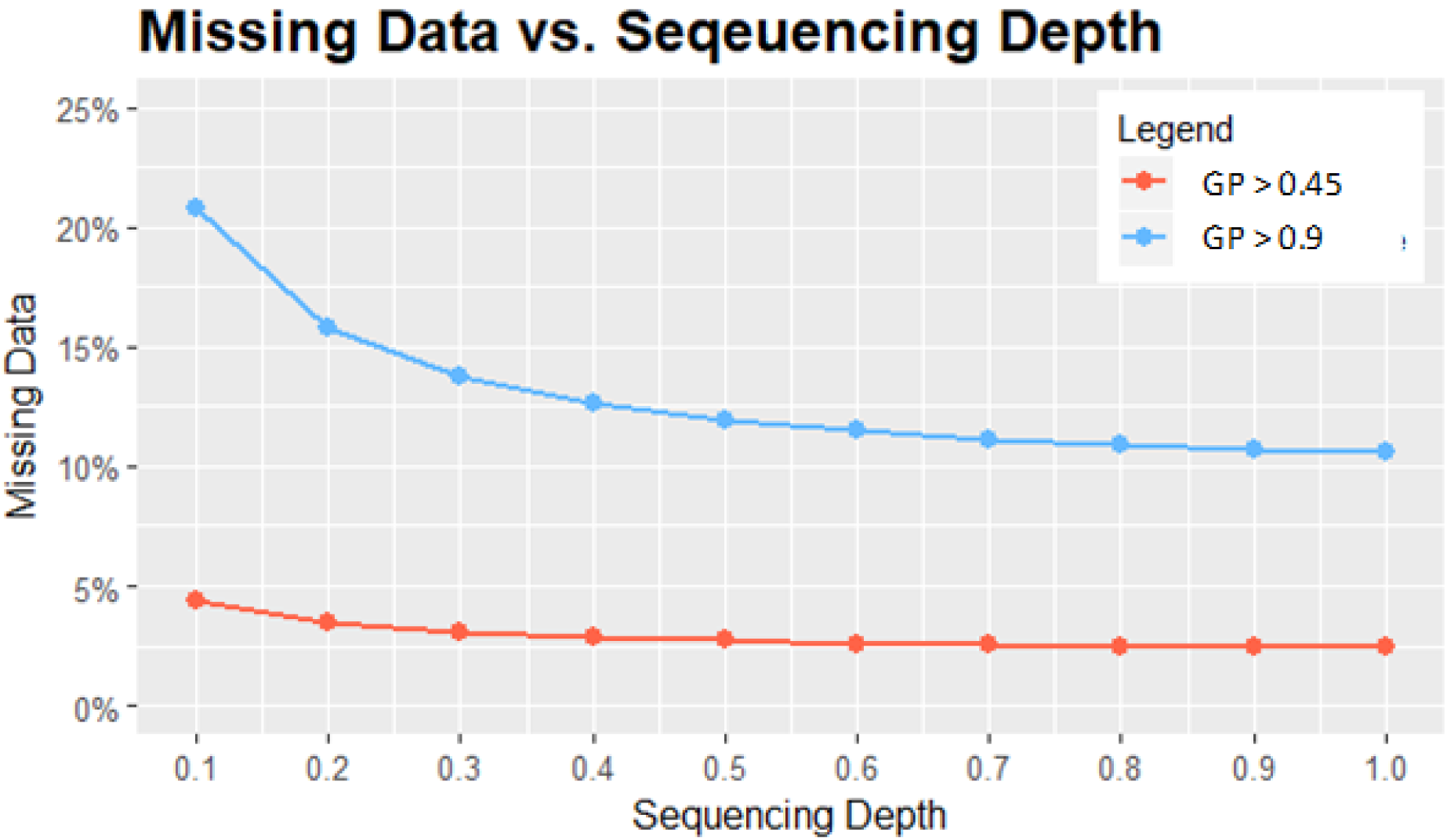
The extent of missing data points reintroduced by filtering for Beagle posterior genotype probabilities greater than 0.45 and 0.9. A mild filter of GP > 0.45 keeps missing data below 5%, while missing data greatly inflates to over 20% at 0.1X with a filter of GP > 0.9.

**Supplementary Figure 2:**
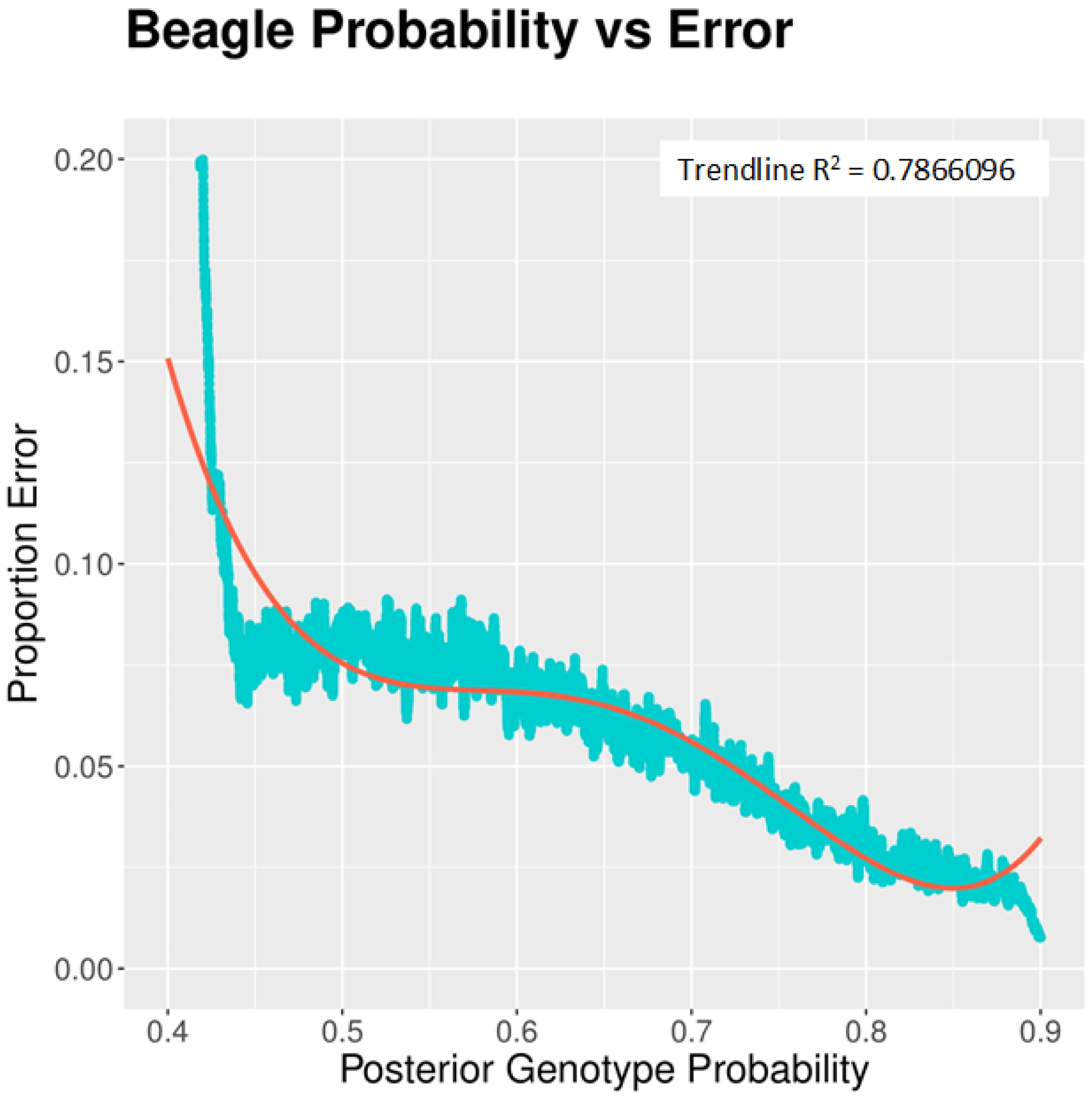
Comparing the frequency of error at individual sites across sequencing depths with the assigned genotype probability for imputed values by Beagle demonstrates a strong correlation, making the posterior genotype probability a useful metric for post imputation filtering for data improvement.

**Supplementary Figure 3:**
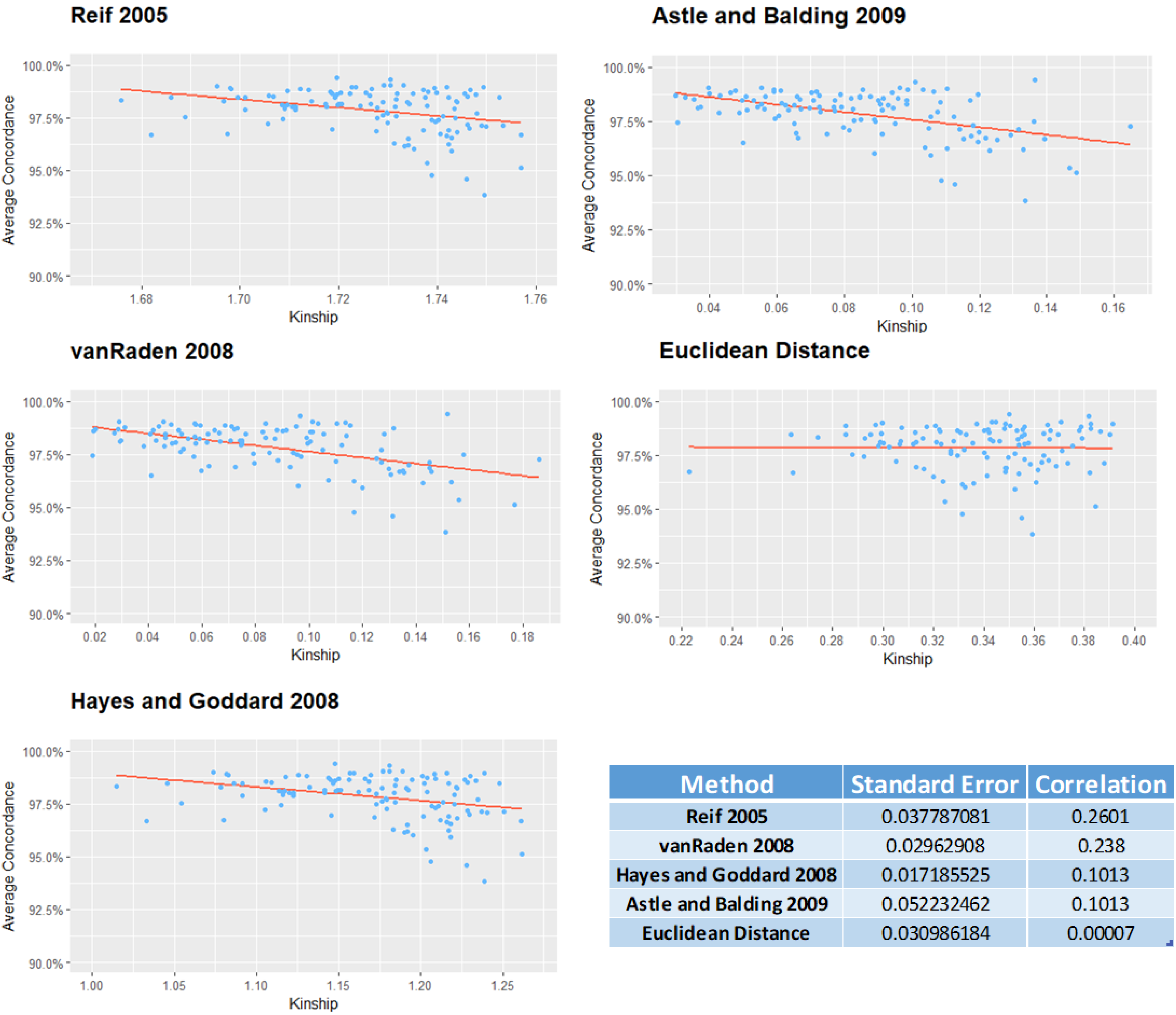
The average of the top five scores for five relatedness metrics are plotted against the average concordance across depths for that genotype. Correlations and standard errors reported in the bottom right hand corner reveal no strong relationship for any metric, which may be explained by the low level variation within our study panel in terms of degree of kinship to the reference panel and overall weak kinship.

**Supplementary Figure 4:**
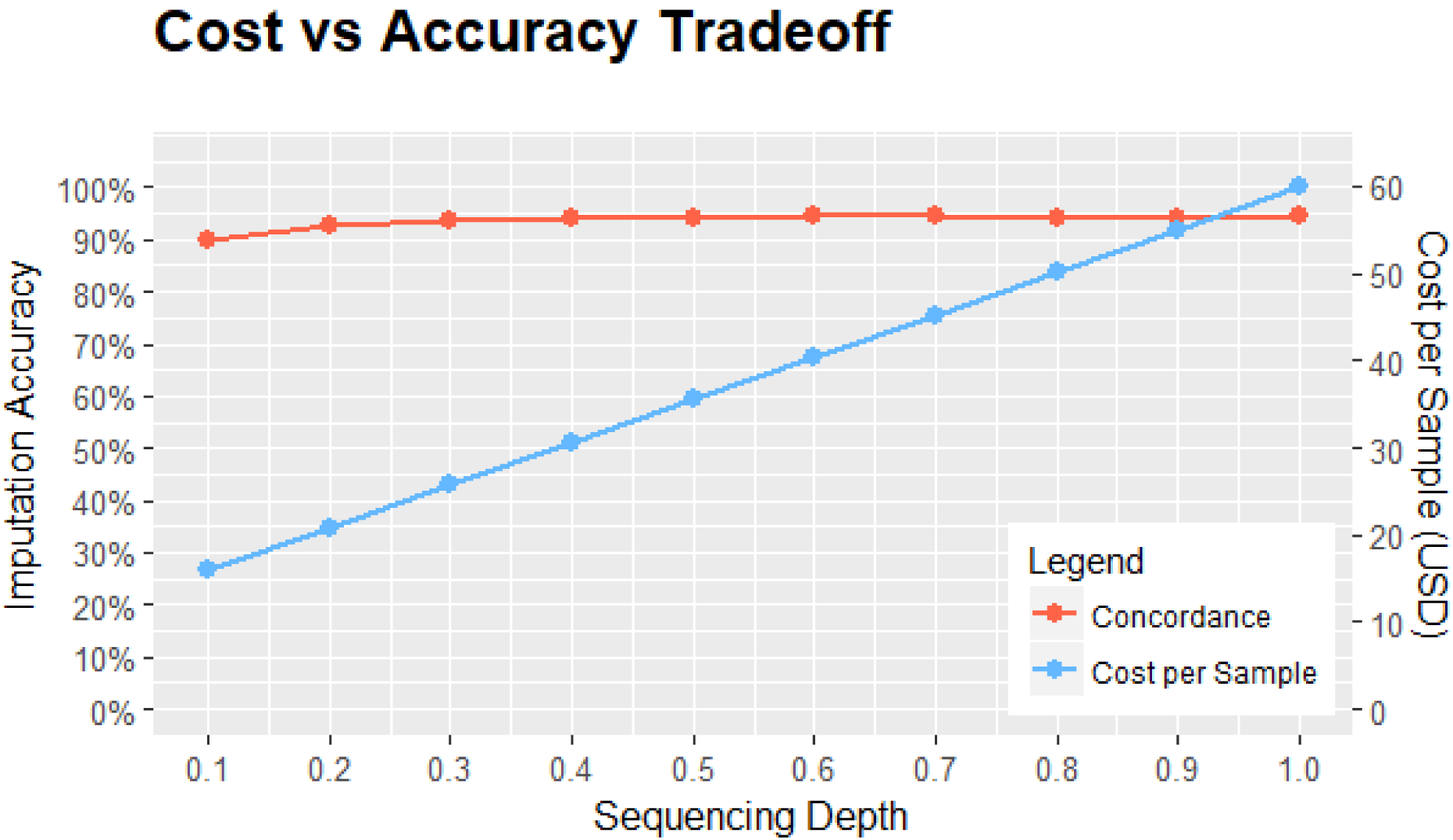
The proportion of accuracy retained relative to 1X from raw imputed values vs the retained cost per sample. Decreasing coverage from 1X to 0.3X results in a nearly negligible loss in accuracy of 0.85%, while decreasing per sample costs by 57.04%

**Supplemental Table 1:**
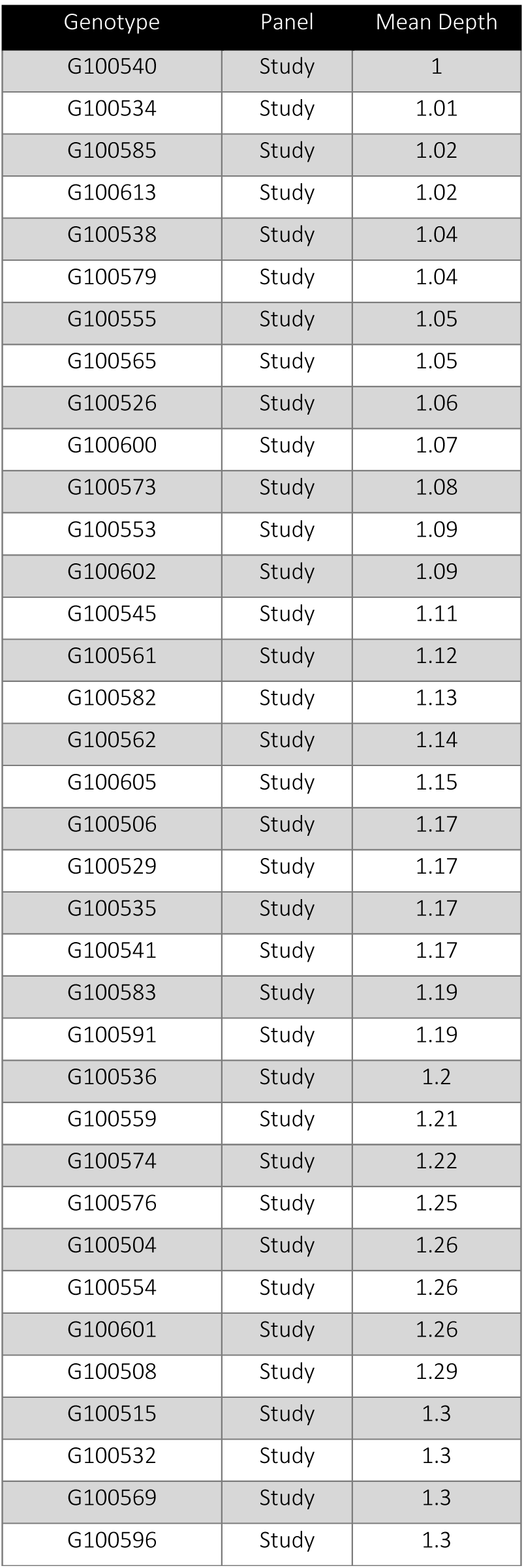
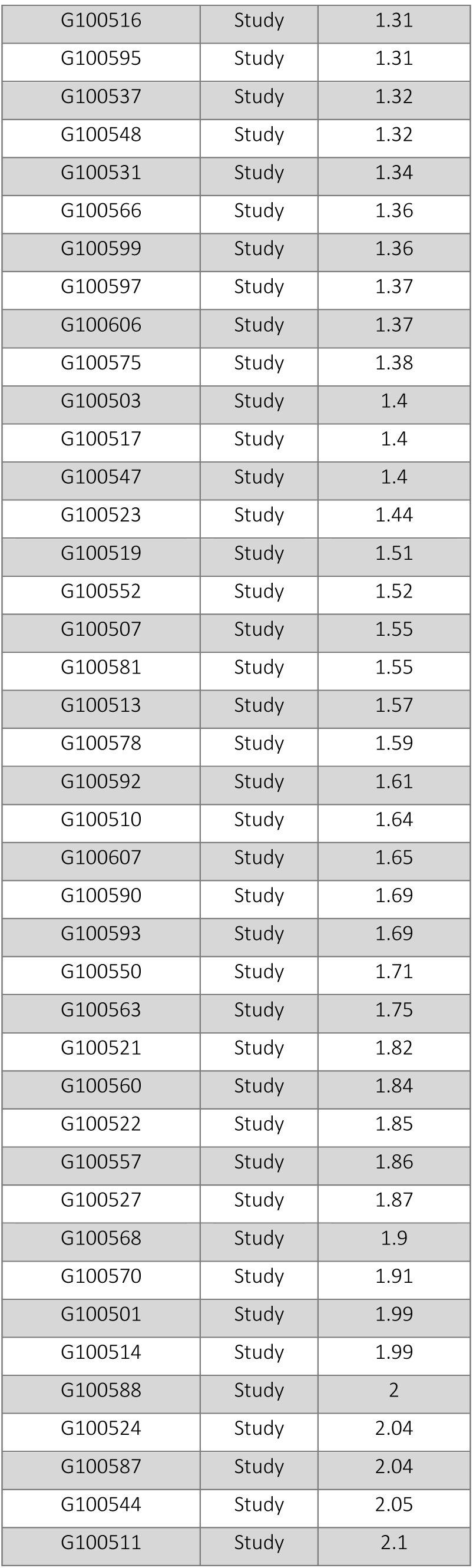
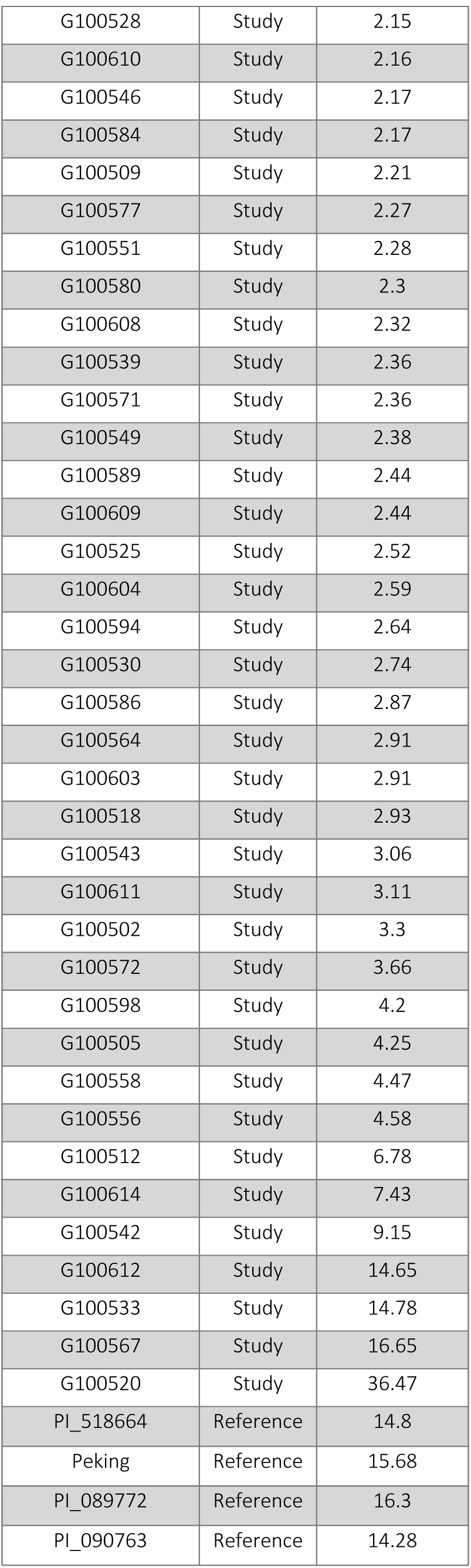
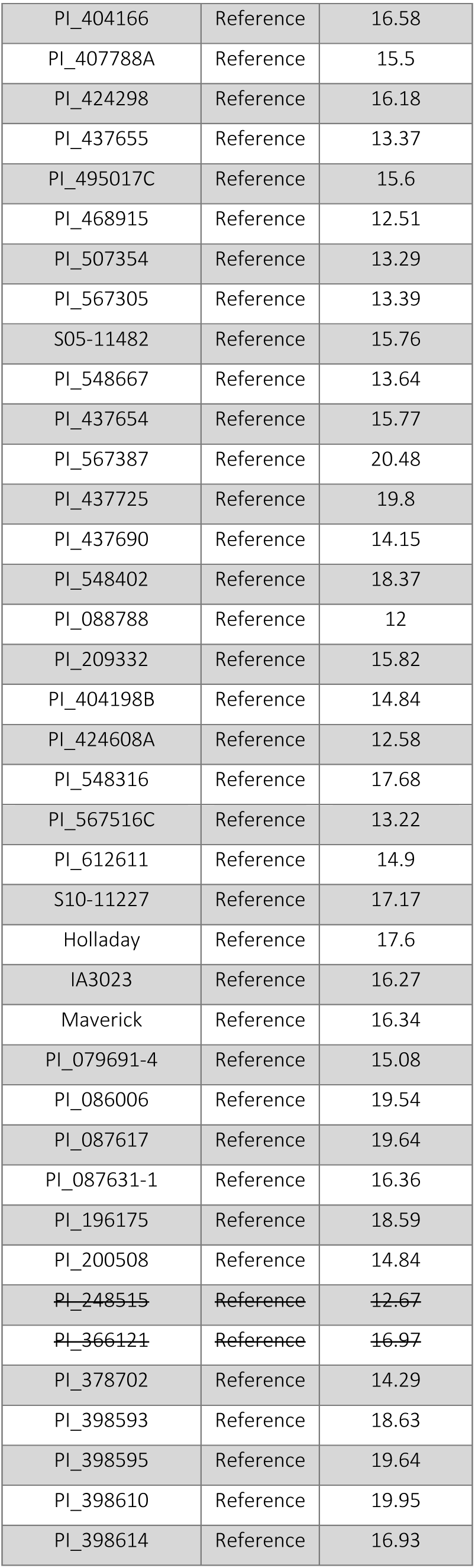
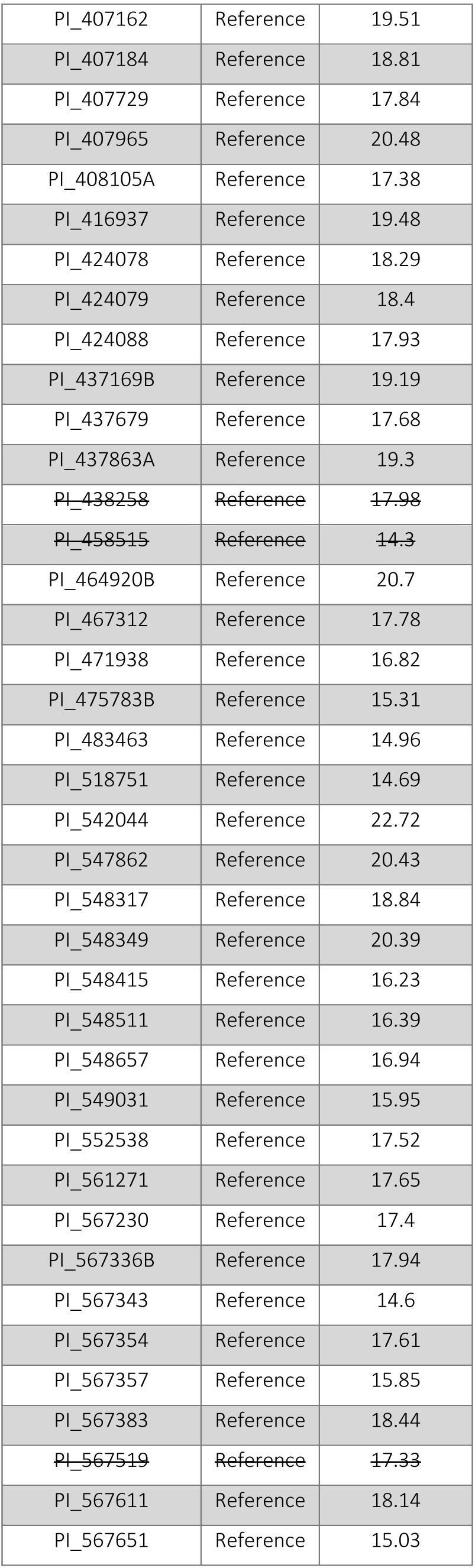
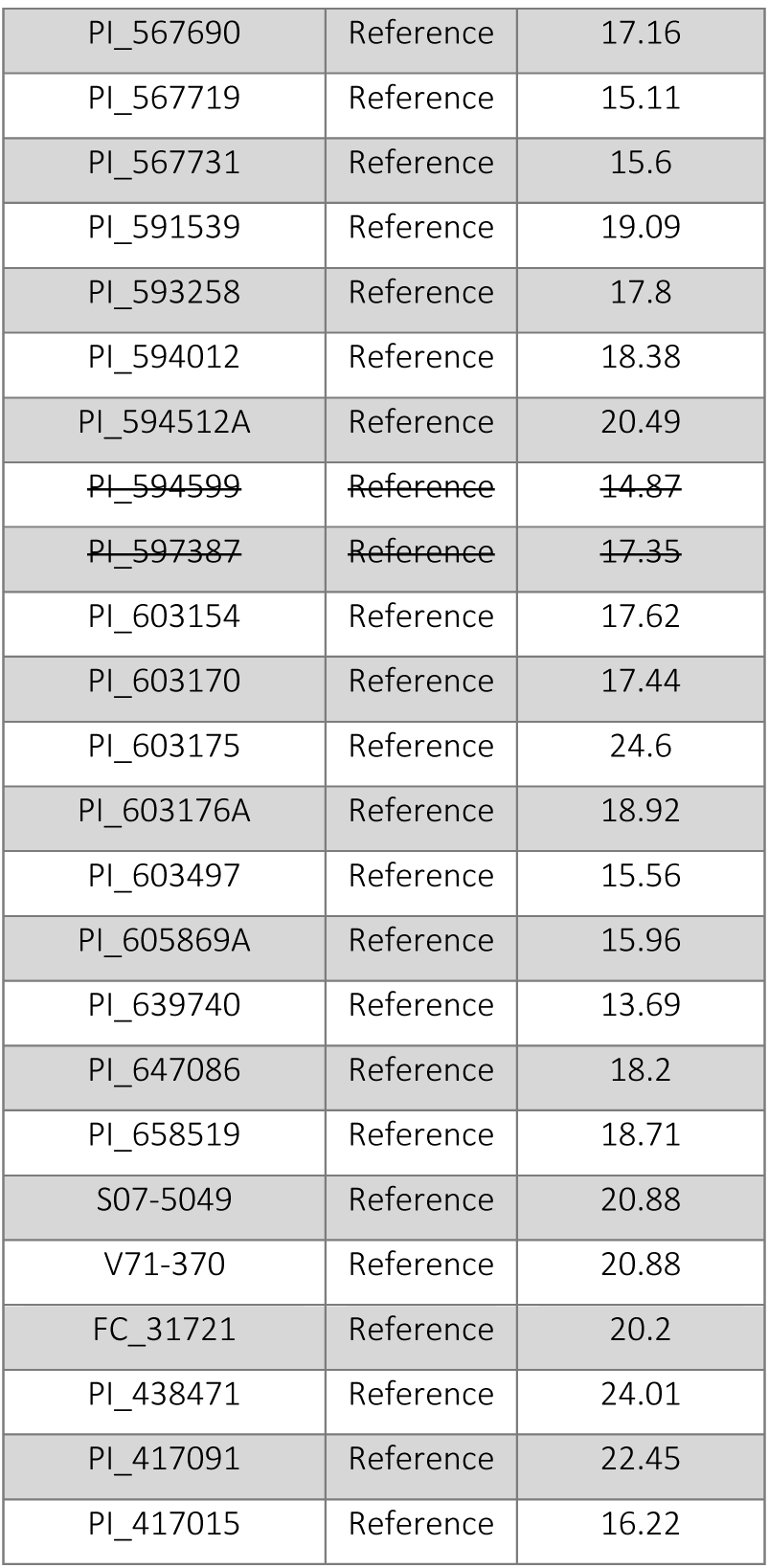
Sequencing depth of individual genotypes used in the generation of the reference and study panels. Samples with text struck through were dropped from final reference panel due to suspected contamination.

| Genotype | Panel | Mean Depth |
| --- | --- | --- |
| G100540 | Study | 1 |
| G100534 | Study | 1.01 |
| G100585 | Study | 1.02 |
| G100613 | Study | 1.02 |
| G100538 | Study | 1.04 |
| G100579 | Study | 1.04 |
| G100555 | Study | 1.05 |
| G100565 | Study | 1.05 |
| G100526 | Study | 1.06 |
| G100600 | Study | 1.07 |
| G100573 | Study | 1.08 |
| G100553 | Study | 1.09 |
| G100602 | Study | 1.09 |
| G100545 | Study | 1.11 |
| G100561 | Study | 1.12 |
| G100582 | Study | 1.13 |
| G100562 | Study | 1.14 |
| G100605 | Study | 1.15 |
| G100506 | Study | 1.17 |
| G100529 | Study | 1.17 |
| G100535 | Study | 1.17 |
| G100541 | Study | 1.17 |
| G100583 | Study | 1.19 |
| G100591 | Study | 1.19 |
| G100536 | Study | 1.2 |
| G100559 | Study | 1.21 |
| G100574 | Study | 1.22 |
| G100576 | Study | 1.25 |
| G100504 | Study | 1.26 |
| G100554 | Study | 1.26 |
| G100601 | Study | 1.26 |
| G100508 | Study | 1.29 |
| G100515 | Study | 1.3 |
| G100532 | Study | 1.3 |
| G100569 | Study | 1.3 |
| G100596 | Study | 1.3 |
| G100516 | Study | 1.31 |
| G100595 | Study | 1.31 |
| G100537 | Study | 1.32 |
| G100548 | Study | 1.32 |
| G100531 | Study | 1.34 |
| G100566 | Study | 1.36 |
| G100599 | Study | 1.36 |
| G100597 | Study | 1.37 |
| G100606 | Study | 1.37 |
| G100575 | Study | 1.38 |
| G100503 | Study | 1.4 |
| G100517 | Study | 1.4 |
| G100547 | Study | 1.4 |
| G100523 | Study | 1.44 |
| G100519 | Study | 1.51 |
| G100552 | Study | 1.52 |
| G100507 | Study | 1.55 |
| G100581 | Study | 1.55 |
| G100513 | Study | 1.57 |
| G100578 | Study | 1.59 |
| G100592 | Study | 1.61 |
| G100510 | Study | 1.64 |
| G100607 | Study | 1.65 |
| G100590 | Study | 1.69 |
| G100593 | Study | 1.69 |
| G100550 | Study | 1.71 |
| G100563 | Study | 1.75 |
| G100521 | Study | 1.82 |
| G100560 | Study | 1.84 |
| G100522 | Study | 1.85 |
| G100557 | Study | 1.86 |
| G100527 | Study | 1.87 |
| G100568 | Study | 1.9 |
| G100570 | Study | 1.91 |
| G100501 | Study | 1.99 |
| G100514 | Study | 1.99 |
| G100588 | Study | 2 |
| G100524 | Study | 2.04 |
| G100587 | Study | 2.04 |
| G100544 | Study | 2.05 |
| G100511 | Study | 2.1 |
| G100528 | Study | 2.15 |
| G100610 | Study | 2.16 |
| G100546 | Study | 2.17 |
| G100584 | Study | 2.17 |
| G100509 | Study | 2.21 |
| G100577 | Study | 2.27 |
| G100551 | Study | 2.28 |
| G100580 | Study | 2.3 |
| G100608 | Study | 2.32 |
| G100539 | Study | 2.36 |
| G100571 | Study | 2.36 |
| G100549 | Study | 2.38 |
| G100589 | Study | 2.44 |
| G100609 | Study | 2.44 |
| G100525 | Study | 2.52 |
| G100604 | Study | 2.59 |
| G100594 | Study | 2.64 |
| G100530 | Study | 2.74 |
| G100586 | Study | 2.87 |
| G100564 | Study | 2.91 |
| G100603 | Study | 2.91 |
| G100518 | Study | 2.93 |
| G100543 | Study | 3.06 |
| G100611 | Study | 3.11 |
| G100502 | Study | 3.3 |
| G100572 | Study | 3.66 |
| G100598 | Study | 4.2 |
| G100505 | Study | 4.25 |
| G100558 | Study | 4.47 |
| G100556 | Study | 4.58 |
| G100512 | Study | 6.78 |
| G100614 | Study | 7.43 |
| G100542 | Study | 9.15 |
| G100612 | Study | 14.65 |
| G100533 | Study | 14.78 |
| G100567 | Study | 16.65 |
| G100520 | Study | 36.47 |
| PI_518664 | Reference | 14.8 |
| Peking | Reference | 15.68 |
| PI_089772 | Reference | 16.3 |
| PI_090763 | Reference | 14.28 |
| PI_404166 | Reference | 16.58 |
| PI_407788A | Reference | 15.5 |
| PI_424298 | Reference | 16.18 |
| PI_437655 | Reference | 13.37 |
| PI_495017C | Reference | 15.6 |
| PI_468915 | Reference | 12.51 |
| PI_507354 | Reference | 13.29 |
| PI_567305 | Reference | 13.39 |
| S05-11482 | Reference | 15.76 |
| PI_548667 | Reference | 13.64 |
| PI_437654 | Reference | 15.77 |
| PI_567387 | Reference | 20.48 |
| PI_437725 | Reference | 19.8 |
| PI_437690 | Reference | 14.15 |
| PI_548402 | Reference | 18.37 |
| PI_088788 | Reference | 12 |
| PI_209332 | Reference | 15.82 |
| PI_404198B | Reference | 14.84 |
| PI_424608A | Reference | 12.58 |
| PI_548316 | Reference | 17.68 |
| PI_567516C | Reference | 13.22 |
| PI_612611 | Reference | 14.9 |
| S10-11227 | Reference | 17.17 |
| Holladay | Reference | 17.6 |
| IA3023 | Reference | 16.27 |
| Maverick | Reference | 16.34 |
| PI_079691-4 | Reference | 15.08 |
| PI_086006 | Reference | 19.54 |
| PI_087617 | Reference | 19.64 |
| PI_087631-1 | Reference | 16.36 |
| PI_196175 | Reference | 18.59 |
| PI_200508 | Reference | 14.84 |
| ~~PI_248515~~ | ~~Reference~~ | ~~12.67~~ |
| ~~PI_366121~~ | ~~Reference~~ | ~~16.97~~ |
| PI_378702 | Reference | 14.29 |
| PI_398593 | Reference | 18.63 |
| PI_398595 | Reference | 19.64 |
| PI_398610 | Reference | 19.95 |
| PI_398614 | Reference | 16.93 |
| PI_407162 | Reference | 19.51 |
| PI_407184 | Reference | 18.81 |
| PI_407729 | Reference | 17.84 |
| PI_407965 | Reference | 20.48 |
| PI_408105A | Reference | 17.38 |
| PI_416937 | Reference | 19.48 |
| PI_424078 | Reference | 18.29 |
| PI_424079 | Reference | 18.4 |
| PI_424088 | Reference | 17.93 |
| PI_437169B | Reference | 19.19 |
| PI_437679 | Reference | 17.68 |
| PI_437863A | Reference | 19.3 |
| ~~PI_438258~~ | ~~Reference~~ | ~~17.98~~ |
| ~~PI_458515~~ | ~~Reference~~ | ~~14.3~~ |
| PI_464920B | Reference | 20.7 |
| PI_467312 | Reference | 17.78 |
| PI_471938 | Reference | 16.82 |
| PI_475783B | Reference | 15.31 |
| PI_483463 | Reference | 14.96 |
| PI_518751 | Reference | 14.69 |
| PI_542044 | Reference | 22.72 |
| PI_547862 | Reference | 20.43 |
| PI_548317 | Reference | 18.84 |
| PI_548349 | Reference | 20.39 |
| PI_548415 | Reference | 16.23 |
| PI_548511 | Reference | 16.39 |
| PI_548657 | Reference | 16.94 |
| PI_549031 | Reference | 15.95 |
| PI_552538 | Reference | 17.52 |
| PI_561271 | Reference | 17.65 |
| PI_567230 | Reference | 17.4 |
| PI_567336B | Reference | 17.94 |
| PI_567343 | Reference | 14.6 |
| PI_567354 | Reference | 17.61 |
| PI_567357 | Reference | 15.85 |
| PI_567383 | Reference | 18.44 |
| ~~PI_567519~~ | ~~Reference~~ | ~~17.33~~ |
| PI_567611 | Reference | 18.14 |
| PI_567651 | Reference | 15.03 |
| PI_567690 | Reference | 17.16 |
| PI_567719 | Reference | 15.11 |
| PI_567731 | Reference | 15.6 |
| PI_591539 | Reference | 19.09 |
| PI_593258 | Reference | 17.8 |
| PI_594012 | Reference | 18.38 |
| PI_594512A | Reference | 20.49 |
| ~~PI_594599~~ | ~~Reference~~ | ~~14.87~~ |
| ~~PI_597387~~ | ~~Reference~~ | ~~17.35~~ |
| PI_603154 | Reference | 17.62 |
| PI_603170 | Reference | 17.44 |
| PI_603175 | Reference | 24.6 |
| PI_603176A | Reference | 18.92 |
| PI_603497 | Reference | 15.56 |
| PI_605869A | Reference | 15.96 |
| PI_639740 | Reference | 13.69 |
| PI_647086 | Reference | 18.2 |
| PI_658519 | Reference | 18.71 |
| S07-5049 | Reference | 20.88 |
| V71-370 | Reference | 20.88 |
| FC_31721 | Reference | 20.2 |
| PI_438471 | Reference | 24.01 |
| PI_417091 | Reference | 22.45 |
| PI_417015 | Reference | 16.22 |

**Supplemental Table 2:** Accuracy improvement as a results of filtering on Beagle’s genotype posterior probability after imputation.

| Depth | Unfiltered Accuracy | GP > 0.45 Accuracy | Improvement | GP > 0.9 Accuracy | Improvement |
| --- | --- | --- | --- | --- | --- |
| 0.1 | 89.70% | 92.70% | 3.00% | 94.80% | 5.10% |
| 0.2 | 92.80% | 95.60% | 2.80% | 97.20% | 4.40% |
| 0.3 | 93.60% | 96.30% | 2.70% | 97.80% | 4.20% |
| 0.4 | 94.20% | 96.60% | 2.40% | 98.10% | 3.90% |
| 0.5 | 94.20% | 96.70% | 2.50% | 98.40% | 4.20% |
| 0.6 | 94.40% | 96.70% | 2.30% | 98.40% | 4.00% |
| 0.7 | 94.50% | 96.70% | 2.20% | 98.50% | 4.00% |
| 0.8 | 94.30% | 96.70% | 2.40% | 98.60% | 4.30% |
| 0.9 | 94.30% | 96.70% | 2.40% | 98.60% | 4.30% |
| 1 | 94.40% | 96.70% | 2.30% | 98.60% | 4.20% |
| **Averages** | **93.64%** | **96.14%** | **2.50%** | **97.90%** | **4.26%** |

**Supplementary Table 3:**
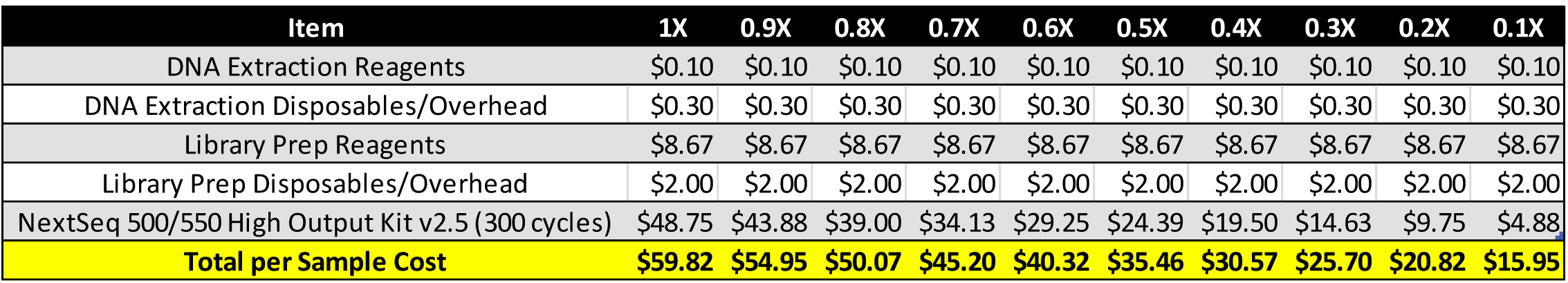
A breakdown of the per item costs involved in the DNA extraction, sample library preparation, and sequencing of genotypes from 0.1X − 1X sequencing depths in USD.

